# Longitudinal structural change of gray matter before and after hemispherotomy

**DOI:** 10.1101/2022.07.07.499164

**Authors:** Hao Yu, Yijun Chen, Junhao Luo, Qingzhu Liu, Peipei Qin, Changtong Wang, Jingli Qu, Lixin Cai, Gaolang Gong

## Abstract

**Background:** Hemispherotomy is an effective surgery developed to treat refractory epilepsy caused by diffuse unihemispheric pathologies. Post-surgery neuroplastic mechanisms supporting behavioral recovery after left and right hemispherotomy as well as their difference remain largely unclear.

**Methods:** In a large cohort of 57 pediatric patients who underwent hemispherotomy, voxel-wise GMV in unaffected regions (contralesional cerebrum and bilateral cerebellum) and behavioral abilities were assessed before and after surgery. Using linear mixed models, we evaluated changes in GMV and behavioral scores, and how GMV changes correlated with improvements in behavioral scores. In addition, three machine learning regression algorithms were applied to assess whether preoperative GMV can predict postoperative behavior.

**Results:** In the two patient groups (29 left hemispherotomy patients, age of surgery mean ± standard deviation = 3.5 ± 2.5; 28 right hemispherotomy patients, age of surgery 4.6 ± 2.5), widespread increases in the postoperative GMV in the contralateral cerebrum and ipsilateral cerebellum but decreases in the postoperative GMV in the contralateral cerebellum were consistently observed. Importantly, the decrease in GMV in the contralateral cerebellum was significantly correlated with improvement in behavioral scores in right but not left hemispherotomy patients. Moreover, the preoperative GMV around the most longitudinally changed locations significantly predicted postoperative behavioral scores in left but not right hemispherotomy patients.

**Conclusions:** Our findings indicate an important role for the contralateral cerebellum in the recovery after right hemispherotomy, and postoperative behavioral scores can be predicted with preoperative GMV features.

*What is already known on this topic:* The contralesional cerebrum plays a key role in the recovery after hemispherotomy.

*What this study adds:* Despite suffering from gray matter atrophy, GMV decrease in contralesional cerebellum is associated with improvement in behavioral score in patients after hemispherotomy.

*How this study might affect research, practice or policy:* These results provide novel insight into the prominence of the contralateral cerebellum in the recovery after hemispherotomy and highlight the clinical role of presurgery GMV in the prediction of postsurgery behavior.

## Introduction

Hemispherotomy can successfully stop seizures in carefully selected pediatric epilepsy patients by disconnecting and disabling the entire affected cerebral hemisphere.^1–4^ In addition to the seize freedom outcome, a number of pediatric patients following such dramatic neurosurgery showed recovery to some degree in certain behavioral domains, ^2–5^ and such recovery has to be underpinned by neuroplastic reorganization in the unaffected brain regions (contralesional cerebrum and bilateral cerebellum).

There have been reported neuroplastic changes following hemispherotomy. For example, animal studies have revealed that the cortex of unaffected hemisphere may project to multiple nuclei and limbs that were normally targeted by the removed cortex. ^6,7^ Increased cortical thickness and metabolism in the unaffected hemisphere was also reported in animals after the surgery.^8,9^ In human, functional neuroimaging studies of hemispherotomy patients suggest that cortical representation originally localized in the disconnected hemisphere can shift to the contralateral hemisphere, accompanied by a certain degree of recovery. ^10–12^ Such shift however has been reported in both preoperative and postoperative period,^11,13^ highlighting the necessity of longitudinal pre- and postoperative data for such investigation. Due to the difficulty of data acquisition in practice, there have been very few longitudinal studies on hemispherectomy or hemispherotomy, and they included either a single patient ^10,12–15^ or 2~10 patients.^11,16,17^ Particularly, these studies focused on neuroplastic changes in functional activation or white matter integrity, leaving structural changes in gray matter unexplored.

Notably, left and right hemisphere damages tend to cause differential functional deficits, e.g., language deficits from left hemisphere damage and hemispatial neglect from right hemisphere damage.^18,19^ Moreover, recent studies further revealed unique structural neuroplastic patterns supporting specific post-damage behavioral recovery between patients of left and right hemisphere damage.^20^ Therefore, it is possible that left and right hemispherotomy are followed by differential post-surgery neuroplastic changes or the same changes but distinct relationships with behavioral recovery.

The present study examined pre- and postsurgery MRI data from two groups of patients who underwent left and right hemispherotomy, aiming to investigate 1) longitudinal changes in gray matter volume (GMV) after surgery in left and right hemispherotomy patients, respectively; 2) whether and how longitudinal GMV changes of the affected regions relate to behavioral recovery within each patient group; and 3) whether presurgery GMV features can predict postsurgery behavioral scores within each patient group.

## Methods

### Patients

Pediatric patients with refractory epilepsy undergoing peri-insular hemispherotomy were included in our present study. Peri-insular hemispherotomy is a surgical variant of functional hemispherotomy in which the affected hemisphere is completely disconnected while resecting a minimal amount of cortical structures to avoid long-term complications.^1^ The inclusion criteria were as follows: 1) underwent peri-insular hemispherotomy between 2016 and 2021 at the Pediatric Epilepsy Center, Peking University First Hospital; 2) at least half a year old when the surgery was performed; 3) available pre- and postoperative MRI images. Initially, 74 patients met these criteria, but the image processing procedures failed for 17 of them due to the poor quality of either the pre- or postoperative MRI images (e.g., severe head motion, bad intensity contrast, abnormalities and severe distortion of the unaffected regions). After excluding these patients, we included 29 pediatric patients who underwent left hemispherotomy and 28 who underwent right hemispherotomy. Demographic and clinical information for the included patients is listed in Table 1. The study was approved by the local Institutional Review Board. Written informed consent was obtained from the legal guardians of the patients.

**Table 1.**
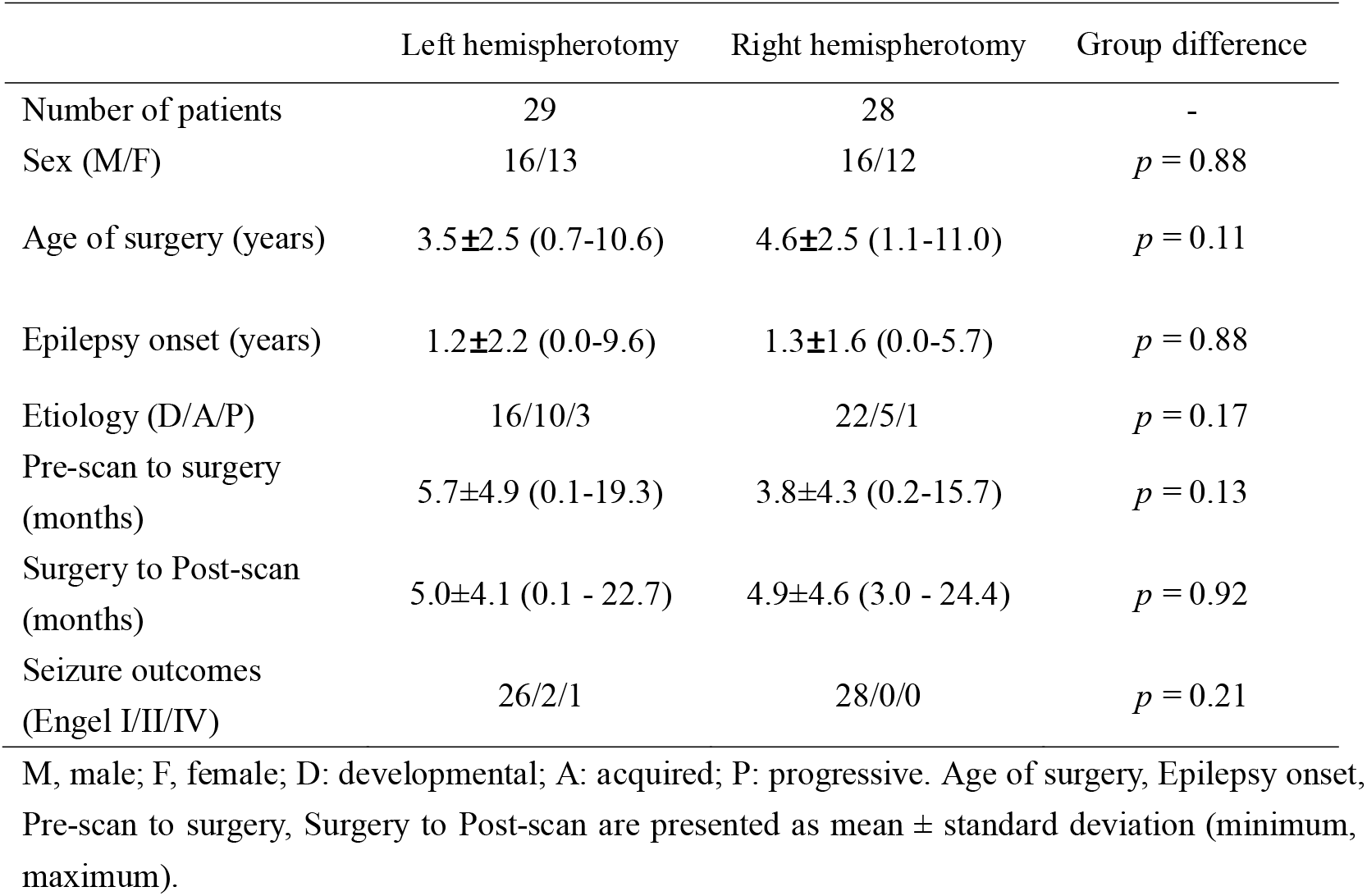
Demographic and clinical data of the included pediatric patients in the present study.

|  | Left hemispherotomy | Right hemispherotomy | Group difference |
| --- | --- | --- | --- |
| Number of patients | 29 | 28 | - |
| Sex (M/F) | 16/13 | 16/12 | $p = 0.88$ |
| Age of surgery (years) | 3.5±2.5 (0.7-10.6) | 4.6±2.5 (1.1-11.0) | $p = 0.11$ |
| Epilepsy onset (years) | 1.2±2.2 (0.0-9.6) | 1.3±1.6 (0.0-5.7) | $p = 0.88$ |
| Etiology (D/A/P) | 16/10/3 | 22/5/1 | $p = 0.17$ |
| Pre-scan to surgery (months) | 5.7±4.9 (0.1-19.3) | 3.8±4.3 (0.2-15.7) | $p = 0.13$ |
| Surgery to Post-scan (months) | 5.0±4.1 (0.1 - 22.7) | 4.9±4.6 (3.0 - 24.4) | $p = 0.92$ |
| Seizure outcomes (Engel I/II/IV) | 26/2/1 | 28/0/0 | $p = 0.21$ |

M, male; F, female; D: developmental; A: acquired; P: progressive. Age of surgery, Epilepsy onset, Pre-scan to surgery, Surgery to Post-scan are presented as mean ± standard deviation (minimum, maximum).

### MRI acquisition

High-quality MRI scanning was performed before and after the surgery (the intervals are listed in Table 1) using a Philips Achieva 3T scanner at the Peking University First Hospital. All T1-weighted scans used the same MPRAGE sequence with the following parameters: axial acquisition; time of repetition (TR) = 8 ms; time of echo (TE) = 3.7 ms; flip angle = 8°; acquisition matrix = 220 × 220, slice thickness = 1 mm; and voxel size = 0.86 × 0.86 × 1.0 mm.

### Behavioral evaluation

The Griffiths Scales of Child Development (3rd edition, GMDS-3) was applied to assess the patient’s behavioral abilities.^21^ The GMDS-3 is a widely used tool for measuring the developmental rate of infants and young children. We examined five domains: foundations of learning, language and communication, eye and hand coordination, personal–social–emotional ability, and gross motor function. The evaluation was usually performed on the same day as the MRI scanning. Only 19 left and 19 right hemispherotomy patients underwent this evaluation both before and after the surgery. Notably, the analysis showed a high correlation across the five domains (mean correlation coefficient reached 0.9), therefore, we performed a factorial analysis using the principal component solution on these five variables, yielding one main factor that accounts for 92.4% percent of the total variance. We used this factor as a single overall behavioral score in our behavior-related analyses. To ensure comparability, we estimated the overall behavioral score for the postoperative GMSD-3 evaluation, also using the weights from the factorial analysis of the preoperative GMSD-3 evaluation.

### Voxel-based morphometry (VBM) processing

For each patient, first, we manually outlined the mask of the cerebral hemisphere that underwent surgery on both the pre- and postoperative T1-weighted images. We then carried out VBM analyses using SPM12 (http://www.fil.ion.ucl.ac.uk/spm). To ensure unbiased comparisons between the left and right hemispheres, a symmetric T1 template in MNI space was constructed using healthy subjects.^26^ Longitudinal VBM processing included the following: 1) for each patient, an average image across the longitudinal images was generated using the Serial Longitudinal Registration toolbox;^27^ 2) the patient-specific average images were bias-corrected and segmented into gray matter (GM), white matter (WM), and cerebrospinal fluid (CSF) probability maps. These tissue-segmented images were then registered to the symmetric template in the MNI space, resulting in a participant-specific GM probability map in MNI space; 3) the GM probability maps were then modulated with twofold Jacobian determinants: from the native space of each time point to the participant-average space and from the participant-average space to the MNI space. For each patient, this resulted in a GMV map in MNI space for each time point, in which each voxel’s value represents its corresponding GMV in the native space; and 4) finally, all of these GMV images were smoothed with an 8-mm full-width half maximum Gaussian kernel. During these procedures, all cerebral hemispheres that underwent surgery were masked out.^28^ The resultant images of each step were carefully checked by visual inspection.

Notably, distinguishing between GM and WM boundaries on the T1w images is difficult for babies younger than 6-8 months (due to a reversal of the normal adult contrasts). ^25^ Children after 8 months likely have an adult-like contrast on the T1 image. We carefully examined the T1 image of our patients particularly at low age and excluded the ones with poor contrast or infantile pattern.

### Identifying longitudinal changes in each patient group

We used a linear mixed-effects model (LMEM) to handle the hierarchical nature of the longitudinal data. The LMEM was applied to evaluate the pre- and postoperative changes in both behavioral scores and voxelwise GMV. Specifically, the ‘fitlme’ function in MATLAB was used. In the model, “MRI to surgery” (i.e., the days between surgery and the pre-/post-MRI scan) and other covariates were modeled as fixed effects, “individual identities” were modeled as random effects. The intercept and slope were allowed to vary across individuals. Specific LMEMs were formulated as follows:

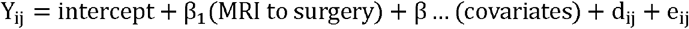

where the intercept and β terms are fixed effects, d_ij_ is the random effect representing within-subject dependence, and e_ij_ represents the residual error.

For each patient group, to determine the pre- and postoperative changes in behavior scores, we applied the LMEM and evaluated the β_1_ term. The onset of epilepsy, age at surgery, sex, and etiology were included as covariates. To evaluate whether and how GMV differed before and after surgery, we performed a voxelwise LMEM analysis within the GM mask of the unaffected regions. During this analysis, the total intracranial volume (ICV) before the surgery was further included as a covariate in the model. Multiple comparisons were corrected using the random field theory (RFT) method (uncorrected *p* < .001), and clusters with a corrected *p* < .05/2 (2 patient groups) were considered significant.

### Correlating longitudinal GMV change with behavioral change

For all identified clusters above, we evaluated whether the GMV change was significantly correlated with the change in overall behavioral score in each patient group. Age at onset of epilepsy, age at surgery, sex, etiology, and TIV were included as covariates.

### Predicting postoperative behavioral scores with preoperative GMV

To evaluate whether preoperative GMV could predict postoperative behavioral scores, we applied three linear machine learning regression algorithms: support vector regression (SVR) (https://www.csie.ntu.edu.tw/~cjlin/libsvm/), relevance vector regression (RVR) (http://www.mlnl.cs.ucl.ac.uk/pronto), and ridge regression (RR) (https://scikit-learn.org). Given the limited sample size, we selected the GMV values from the locations showing the greatest local changes in GMV before and after surgery as input features. Specially, we identified the local peaks within the T map from the LMEM using the AFNI program ‘3dmaxima’ with a threshold value of 4.5. This resulted in 26 local peaks for each group. For each local peak, the mean GMV value was extracted from a sphere of radius of 7.5 mm at the peak coordinate. These locations represent the strongest developmental or plastic regions following surgery, therefore, their preoperative GMV is more likely predictive of postoperative behaviors than that of other regions.

For each algorithm, nested leave-one-out-cross-validation (LOOCV) was applied, with an outer LOOCV loop estimating the generalizability of the model and an inner LOOCV loop determining the optimal parameter (e.g., *C* parameter for SVR) if any. Specifically, N-1 patients were used as the training set for each outer LOOCV loop, where N is the total number of patients, and the remaining patient was used as the testing sample. This procedure was repeated N times, with each patient being the testing sample once. Within each loop of the outer LOOCV, the inner LOOCVs were applied to determine the optimal parameter for the outer LOOCV fold to construct a prediction model, which was then used to predict the score of the testing sample.

To evaluate the performance of the predictive model for each algorithm, the Pearson correlation (*r*) and normalized root mean square error (NRMSE) were calculated. A permutation test (1,000 times) was applied to determine whether the evaluation indicators (r and NRMSE) were significantly better than those expected by chance.

### Data Availability

The data that support the findings of this study are available upon reasonable request from the corresponding author. Data are not publicly available as they include patient data that could compromise the privacy of participants.

## Results

As shown in Table 1, there was no significant difference in age, sex, or onset or etiology of epilepsy between the left and right hemispherotomy groups. Seizure outcome was assessed using the Engel scale, ^26^ and 90% of left and 100% of right hemispherotomy patients reached Engel 1 at the follow-up of 3 months (Chi-square test, p = 0.21).

### Longitudinal behavioral changes in each patient group

For each patient, an overall behavioral score was obtained using the GMDS-3 evaluation. Neither the preoperative nor postoperative behavioral scores showed a significant difference between the two patient groups (two-sample t tests, ps > 0.05). For both groups, the LMEM showed a significant increase in the postoperative score compared with the preoperative behavioral score (left hemispherotomy: t = 2.10, p = 0.04; right hemispherotomy: t = 3.97, p <0.001).

### Longitudinal GMV changes in each patient group

To identify local GMV changes in each patient group before and after the surgery, we applied a voxelwise LMEM search within the GM mask of the unaffected regions. For the two patient groups, the resultant T maps of GMV change overall mirrored each other (after flipping one map, r = 0.87), suggesting similar spatial patterns of ipsilateral and contralateral GMV changes between the two groups (Figure 2).

**Figure 1.**
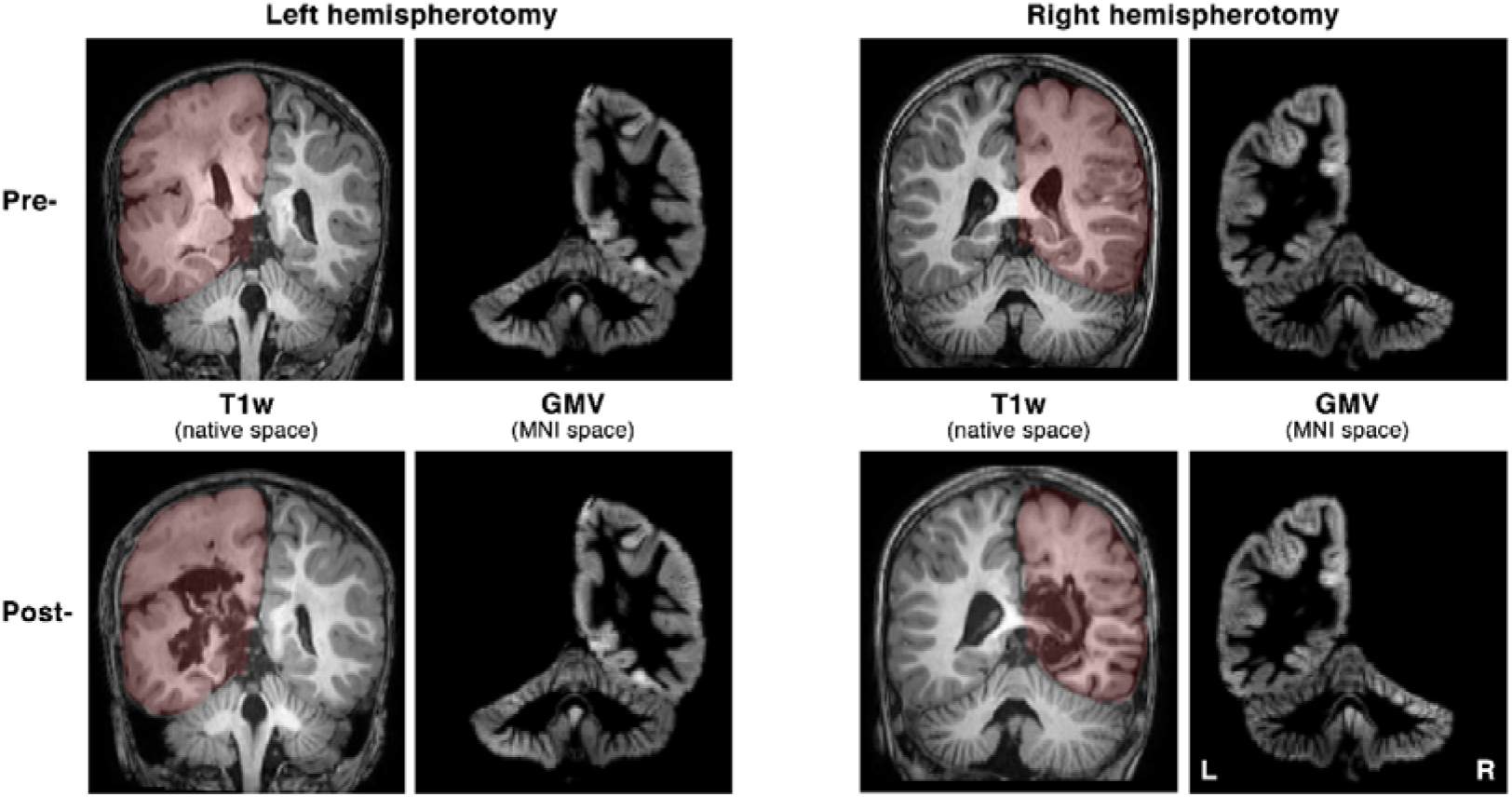
T1-weighted (T1w) and gray matter volume (GMV) images of example left and right hemispherotomy patients. The cerebral hemisphere was manually masked out (indicated by red) in the T1w native space and excluded in the GMV analyses of the MNI space. Pre: Preoperative; Post: Postoperative.

**Figure 2.**
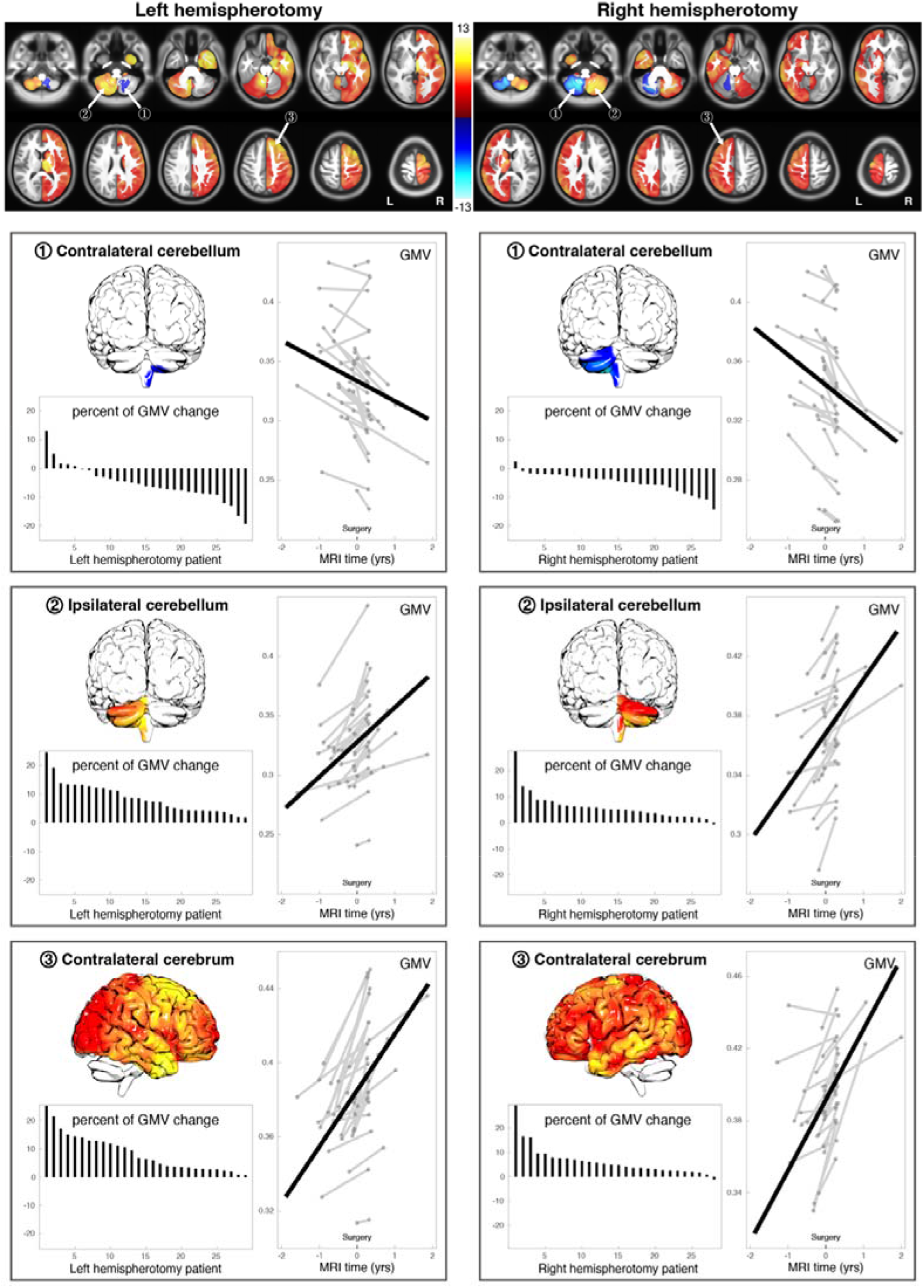
GMV change before and after surgery for the two patient groups. The results for the left and right hemispherotomies are shown in the left and right columns, respectively. The first row represents the T maps of the GMV change from the linear mixed model for the two groups. The significant identified clusters are denoted by arrows. For each cluster (rows 2 to 4), the decrease or increase in percent change in GMV in all patients is illustrated as a bar graph; the fitted longitudinal change in GMV before and after the surgery is represented as a thick black line within the panel.

After correction for multiple comparisons, we observed 3 significant clusters showing GMV changes before and after surgery for either the left or right hemispherotomy group (Figure 2). In both groups, the largest cluster covered almost the entire contralateral cerebrum and exhibited significantly increased GMV after the surgery (left hemispherotomy: t = 7.21, p < 0.001; right hemispherotomy: t = 7.68, p < 0.001). At the individual level, all left and 96.4% of right hemispherotomy patients showed an increase in raw GMV in this cluster. The second-largest cluster was located around the entire ipsilateral cerebellum, which also exhibited significantly increased GMV after the surgery (left hemispherotomy: t = 8.69, p < 0.001; right hemispherotomy: t = 8.67, p < 0.001). All left and right hemispherotomy patients had an increase in raw GMV in this cluster.

In contrast to the two largest clusters, the remaining cluster in the contralateral cerebellum showed significantly decreased GMV after the surgery (left hemispherotomy: t = −4.65, p < 0.001; right hemispherotomy: t = −7.82, p < 0.001), and the cluster size of the left hemispherotomy group was smaller than that of the right hemispherotomy group. At the individual level, 79.5% of the left and 96.4% of right hemispherotomy patients showed a decrease in raw GMV in this cluster.

The percent GMV change in all patients for the three clusters is illustrated in Figure 2 (contralateral cerebrum: left hemispherotomy, mean ± SD = 8.5% ± 6.3%; right hemispherotomy, 6.3% ± 5.9%; ipsilateral cerebellum: left hemispherotomy, 9.0% ± 5.6%; right hemispherotomy, 6.3% ± 5.4%; contralateral cerebellum: left hemispherotomy, −5.2% ± 6.4%; right hemispherotomy, mean −4.9% ± 3.5%). The percent change in GMV was not significantly different between the left and right hemispherotomy patient groups in any of the clusters (t-test, p > 0.05), again supporting the similarity of ipsilateral and contralateral GMV changes between the two groups.

### Correlation between longitudinal GMV and behavioral change

For each patient group, we evaluated the correlation between longitudinal GMV changes in each cluster and behavioral score changes before and after the surgery using the Pearson correlation coefficient. As shown in Figure 3, the change in GMV in the contralateral cerebellum was negatively correlated with the change in overall behavioral score in the right hemispherotomy patients (r = −0.61, p = 0.02; after removing one outliner: r = −0.81, p < 0.001) but not in the left hemispherotomy patients (r = 0.09, p = 0.77). Given the negative direction of the GMV change in the contralateral cerebellum and the positive direction of the behavioral change, the observed negative correlation here indicates that a greater GMV decrease was associated with greater behavioral improvement in right hemispherotomy patients.

**Figure 3.**
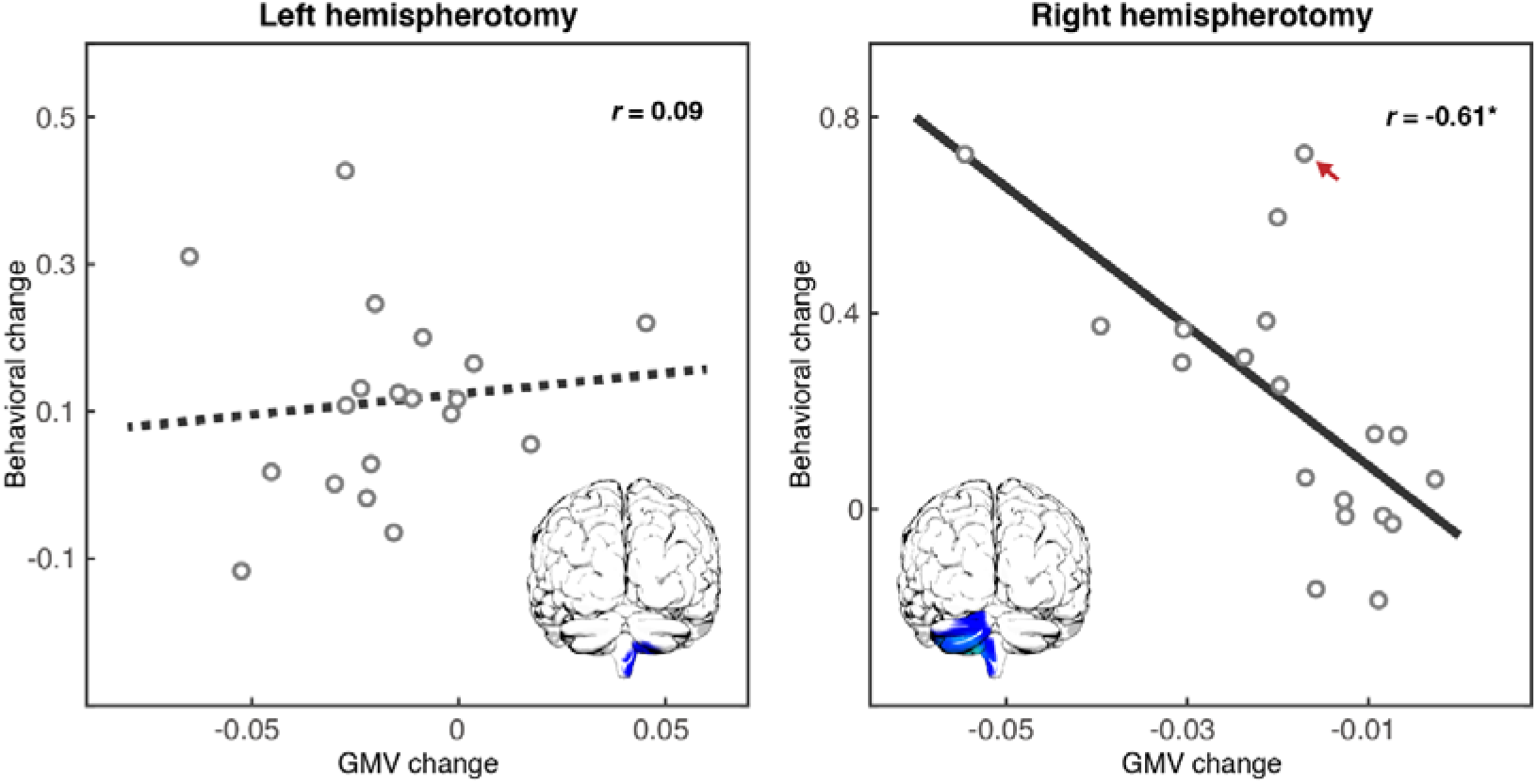
Correlation between the longitudinal change in GMV in the contralateral cerebellum and the change in behavioral score. As indicated by the red arrow in the scatter plot, one outlying datapoint was identified, according to Cook’s distance and leverage value. The correlation reached −0.81 (p < 0.001) when the outlying datapoint was removed.

No significant correlation was found between the GMV increase in the contralateral cerebrum or ipsilateral cerebellum and behavioral change in either group. Given the extensive coverage of the cluster of the contralateral cerebrum, we further did a voxel-wise analysis searching for GMV-behavior correlations across the contralateral cerebrum within both groups, but found no significant results neither.

### Prediction of the postoperative behavioral score with preoperative GMV

For each patient, preoperative GMV values from 26 unaffected GM locations showing local maximum GMV changes after the surgery were selected as predictive features (Figure 4). As shown in Table 2, the preoperative GMV is strongly predictive of postoperative behavioral abilities in left hemispherotomy patients. In contrast, the preoperative overall GMV can not significantly predict the postoperative behavioral scores for right hemispherotomy patients, regardless of the algorithm used.

**Figure 4.**
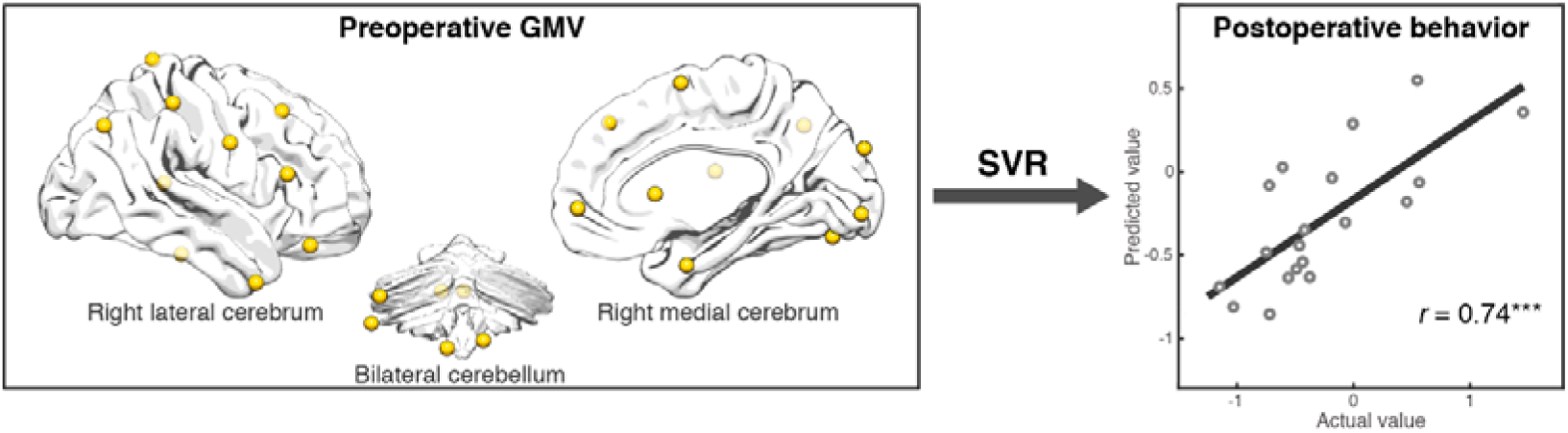
Prediction of postoperative behavior with preoperative GMV features in left hemispherotomy patients. The preoperative GMVs from 26 unaffected GM locations showing local maximum changes in GMV after the surgery (indicated as yellow spheres) were selected as predictive features. SVR: support vector regression.

**Table 2.** Prediction of postoperative behavioral scores with preoperative gray matter volume (GMV) features.

| Predictive model | Left hemispherotomy group |  | Right hemispherotomy group |  |
| --- | --- | --- | --- | --- |
| | $r$ | NRMSE | $r$ | NRMSE |
|  | (p value) | (p value) | (p value) | (p value) |
| SVR | **0.74 (0.001\*)** | **0.16 (0.001\*)** | 0.06 (0.54) | 0.41 (0.18) |
| RVR | **0.66 (0.002\*)** | **0.19 (0.002\*)** | 0.24 (0.10) | 0.30 (0.10) |
| RR | **0.62 (0.003\*)** | **0.20 (0.001\*)** | 0.13 (0.21) | **0.31 (0.003\*)** |

SVR: support vector regression; RVR: relevance vector regression; RR: ridge regression; $r$ : Pearson correlation coefficient for the postoperative behavioral score and its predictive value; NRMSE: normalized root mean square error. Bold represents predictive ability significantly better than chance.

## Discussion

Using both pre- and postsurgery MRI data from pediatric patients who underwent hemispherotomy, we examined GMV changes before and after surgery as well as their relationship with behavioral improvements for the first time. In both left and right hemispherotomy groups, the results consistently showed a widespread increase in postsurgery GMV in the contralateral cerebrum and ipsilateral cerebellum but a decrease in postsurgery GMV in the contralateral cerebellum. Importantly, the decrease in GMV in the contralateral cerebellum was significantly correlated with improvement of behavioral scores in right hemispherotomy patients but not in left hemispherotomy patients. Moreover, the presurgery GMV around the most longitudinally changed locations significantly predicted the postsurgery behavioral scores in left hemispherotomy patients but not in right hemispherotomy patients. These results provide novel insight into morphological change of GM underlying behavioral recovery in hemispherotomy patients and highlight the clinical role of presurgery GMV in the prediction of postsurgery behavior.

Postsurgery GMV changes following hemispherotomy could represent following mechanisms: neurodevelopment, neurodegeneration, and neuroplastic reorganization. While the lack of longitudinal MRI data from age-matched healthy controls leads to difficulties in differentiating between neurodevelopment-induced and surgery-induced changes in GMV, the two patient groups *per se* provide excellent age-matched controls to each other. Since both cerebral and cerebellar GMV showed a progressive increase during healthy early childhood (i.e., the age range of our pediatric patients),^30–32^ our observed GMV increase in the contralateral cerebrum and ipsilateral cerebellum in both patient groups likely represents both normal neurodevelopment and neuroplastic reorganization following hemispherotomy.

Interestingly, the observed longitudinal GMV decrease in the contralateral cerebellum contrasts with the normal neurodevelopmental pattern and therefore is likely related to neuroplastic reorganization or neurodegeneration following surgery. In fact, a critical role of the contralateral cerebellum in post-cerebral hemispherectomy recovery has been demonstrated in rats: contralateral hemicerebellectomy two weeks after cerebral hemispherectomy did not impair functional recovery, but contralateral hemicerebellectomy before or simultaneous with cerebral hemispherectomy led to significantly worse functional recovery.^33^ In line with this, our currently observed contralateral cerebellar GMV decrease did show a positive effect: a greater GMV decrease was associated with better behavioral improvement in our right hemispherotomy patients, which highlights the neuroplastic nature for such GMV decrease. Compatibly, stroke patients also showed the positive role of GMV decrease in functional recovery, e.g., greater decrease of cerebellar GMV, greater improvement of post-stroke Barthel index^27^; greater decrease of GMV around middle cingulate cortex, greater improvement of post-stroke motor recovery^20^. However, it is possible that crossed cerebellar diaschisis and progressive neurodegeneration also contributes to our observed GMV decrease to some degree.

Predicting postsurgery functional or behavioral status preoperatively is crucial for children undergoing hemispherotomy. A set of preoperative MRI-based parameters has been reported to predict postsurgery behavioral outcomes, e.g., contralateral MRI abnormalities,^35,36^ asymmetries in the brain stem or corticospinal tracts within the brain stem^7,34,37–41^, and sensorimotor fMRI results. ^39^ The results of the present study highlight the predictive value of preoperative GMV features, and they can easily be derived from routine structural MRI of hemispherotomy patients. Therefore, clinically applicable prediction tools with preoperative GMV features could be developed for various post-cerebral hemispherotomy outcomes. Importantly, the present study proposed a multivariate machine learning-based framework for postoperative prediction, complementary to previous univariate-based prediction models. This multivariate framework showed robust results and can easily be applied for the prediction of other postoperative outcomes. The GMV feature selection in our current framework was implemented using previously obtained information (i.e., the locations that changed the most longitudinally), but sophisticated data-driven approaches are required for practical usage in the future.

Notably, the present study divided left and right hemispherotomy patients into separate groups. Given the hemispheric specialization and asymmetries, we expected common and unique GMV changes after left and right hemispherotomy, as previous studies reported in unilateral stroke patients.^20^ However, the two pediatric patient groups showed a quite mirrored pattern of postsurgery GMV change, likely reflecting the similar neuroplasticity of the two hemispheres and unestablished structural lateralization at an early developmental stage.^39,40^ Despite, similar changes in the two hemispheres may lead to different behavioral consequences, e.g., a brain-behavior correlation in right but not left hemispherotomy and prediction of post-operative behavior in left but not right hemispherotomy. This may relate to the difference of functional contribution and plasticity of each hemisphere following such a dramatic neurosurgery, which warrants further investigation in the future.

Finally, a few limitations should be addressed. The relative sample size of our study was quite large for a longitudinal MRI study of hemispherotomy, but the absolute sample size is limited. More patients are required, particularly for generalizable machine learning-based postoperative prediction. Next, the age range of our patients was relatively large (0.7~11 yrs), and age-related neurodevelopmental differences among patients may have confounded our results to some degree, even after including patient age as a covariate in the statistical model.

## Conclusions

Decreased GMV in the contralateral cerebellum following right hemispherotomy correlates with improvement in behavioral scores, indicating a critical role of contralateral cerebellum in post-surgery recovery. Postoperative behavioral scores can be predicted with preoperative GMV features, highlighting the clinically relevant predictive role of GMV before surgery.

## Acknowledgements

The authors thank Qian Zhang from Peking University First Hospital for her technical assistance.

## References

1. Marras CE, Granata T, Franzini A, et al. Hemispherotomy and functional hemispherectomy: Indications and outcome. Epilepsy Res. 2010;89:104–112.

2. Moosa ANV, Gupta A, Jehi L, et al. Longitudinal seizure outcome and prognostic predictors after hemispherectomy in 170 children. Neurology. 2013;80:253–260.

3. Kossoff EH, Vining EPG, Pillas DJ, et al. Hemispherectomy for intractable unihemispheric epilepsy Etiology vs outcome. Neurology. 2003;61:887–890.

4. Jonas R, Nguyen S, Hu B, et al. Cerebral hemispherectomy: Hospital course, seizure, developmental, language, and motor outcomes. Neurology. 2004;62:1712–1721.

5. Moosa ANV, Jehi L, Marashly A, et al. Long-term functional outcomes and their predictors after hemispherectomy in 115 children. Epilepsia. 2013;54:1771–1779.

6. Gómez-Pinilla F, Villablanca JR, Sonnier BJ, Levine MS. Reorganization of Pericruciate cortical projections to the spinal cord and dorsal column nuclei after neonatal or adult cerebral hemispherectomy in cats. Brain Res. 1986;385:343–355.

7. Machado AGG, Shoji A, Ballester G, Marino R. Mapping of the Rat’s Motor Area after Hemispherectomy: The Hemispheres as Potentially Independent Motor Brains. Epilepsia. 2003;44:500–506.

8. Hovda DA, Villabianca JR, Chugani HT, Phelps ME. Cerebral metabolism following neonatal or adult hemineodecortication in cats: I. Effects on glucose metabolism using [14C] 2-deoxy-D-glucose autoradiography. J Cereb Blood Flow Metab. SAGE Publications Sage UK: London, England; 1996;16:134–146.

9. Schmanke TD, Villablanca JR. Regional Age-Dependent Effects of Hemineodecortication upon Contralateral Neocortical Thickness: Comparison with Other Measures of Cortical Size. Dev Neurosci. 1999;21:290–297.

10. Pilato F, Dileone M, Capone F, et al. Unaffected motor cortex remodeling after hemispherectomy in an epileptic cerebral palsy patient. A TMS and fMRI study. Epilepsy Res. 2009;85:243–251.

11. Zhang J, Mei S, Liu Q, et al. fMRI and DTI assessment of patients undergoing radical epilepsy surgery. Epilepsy Res. 2013;104:253–263.

12. Hertz□Pannier L, Chiron C, Jambaqué I, et al. Late plasticity for language in a child’s non□dominant hemisphere: A pre□ and post□surgery fMRI study. Brain. 2002;125:361–372.

13. Rutten G-JM, Ramsey NF, Van Rijen PC, Franssen H, Van Veelen CWM. Interhemispheric Reorganization of Motor Hand Function to the Primary Motor Cortex Predicted With Functional Magnetic Resonance Imaging and Transcranial Magnetic Stimulation. J Child Neurol. 2002;17:292–297.

14. Chiron C, Raynaud C, Jambaqué I, Dulac O, Zilbovicius M, Syrota A. A serial study of regional cerebral blood flow before and after hemispherectomy in a child. Epilepsy Res. 1991;8:232–240.

15. Danelli L, Cossu G, Berlingeri M, Bottini G, Sberna M, Paulesu E. Is a lone right hemisphere enough? Neurolinguistic architecture in a case with a very early left hemispherectomy. Neurocase. 2013;19:209–231.

16. Govindan RM, Brescoll J, Chugani HT. Cerebellar Pathway Changes Following Cerebral Hemispherectomy. J Child Neurol. 2013;28:1548–1554.

17. Rosazza C, Deleo F, D’Incerti L, et al. Tracking the Re-organization of Motor Functions After Disconnective Surgery: A Longitudinal fMRI and DTI Study. Front Neurol. 2018;9:400.

18. Price CJ. The anatomy of language: contributions from functional neuroimaging. J Anat. Cambridge University Press; 2000;197:335–359.

19. Corbetta M, Shulman GL. Spatial neglect and attention networks. Annu Rev Neurosci. Annual Reviews; 2011;34:569–599.

20. Chen Y, Jiang Y, Kong X, et al. Common and unique structural plasticity after left and right hemisphere stroke. J Cereb Blood Flow Metab. Epub 2021 Aug 20.:0271678X2110366.

21. Luiz DM, Foxcroft CD, Povey J-L. The Griffiths Scales of Mental Development: A Factorial Validity Study. South Afr J Psychol. 2006;36:192–214.

22. Brett M, Leff AP, Rorden C, Ashburner J. Spatial Normalization of Brain Images with Focal Lesions Using Cost Function Masking. NeuroImage. 2001;14:486–500.

23. Ashburner J, Ridgway GR. Symmetric diffeomorphic modeling of longitudinal structural MRI. Front Neurosci. 2013;6:197.

24. Crinion J, Ashburner J, Leff A, Brett M, Price C, Friston K. Spatial normalization of lesioned brains: Performance evaluation and impact on fMRI analyses. NeuroImage. 2007;37:866–875.

25. Dubois J, Dehaene-Lambertz G, Kulikova S, Poupon C, Hüppi PS, Hertz-Pannier L. The early development of brain white matter: A review of imaging studies in fetuses, newborns and infants. Neuroscience. 2014;276:48–71.

26. Engel Jr J. Outcome with respect to epileptic seizures. Surg Treat Epilepsies. Raven Press; Epub 1993.:609–621.

27. Yu X, Yang L, Song R, et al. Changes in structure and perfusion of grey matter tissues during recovery from Ischaemic subcortical stroke: a longitudinal MRI study. Eur J Neurosci. 2017;46:2308–2314.

28. Sprague JM. Interaction of cortex and superior colliculus in mediation of visually guided behavior in the cat. Science. American Association for the Advancement of Science; 1966;153:1544–1547.

29. Marino, Jr. R, Machado AGG, Timo-Iaria C. Functional Recovery after Combined Cerebral and Cerebellar Hemispherectomy in the Rat. Stereotact Funct Neurosurg. 2001;76:83–93.

30. Boshuisen K, van Schooneveld MMJ, Leijten FSS, et al. Contralateral MRI abnormalities affect seizure and cognitive outcome after hemispherectomy. Neurology. 2010;75:1623–1630.

31. Hallbook T, Ruggieri P, Adina C, et al. Contralateral MRI abnormalities in candidates for hemispherectomy for refractory epilepsy. Epilepsia. 2010;51:556–563.

32. Wakamoto H, Eluvathingal TJ, Makki M, Juhasz C, Chugani HT. Diffusion Tensor Imaging of the Corticospinal Tract Following Cerebral Hemispherectomy. J Child Neurol. 2006;21:566–571.

33. Mullin JP, Soni P, Lee S, et al. Volumetric Analysis of Cerebral Peduncles and Cerebellar Hemispheres for Predicting Hemiparesis After Hemispherectomy: Neurosurgery. 2016;79:499–507.

34. Nelles M, Urbach H, Sassen R, et al. Functional hemispherectomy: postoperative motor state and correlation to preoperative DTI. Neuroradiology. 2015;57:1093–1102.

35. Du X-Y, Chen S-C, Guan Y-G, et al. Asymmetry of Cerebral Peduncles for Predicting Motor Function Restoration in Young Patients Before Hemispherectomy. World Neurosurg. 2018;116:e634–e639.

36. Wang AC, Ibrahim GM, Poliakov AV, et al. Corticospinal tract atrophy and motor fMRI predict motor preservation after functional cerebral hemispherectomy. J Neurosurg Pediatr. 2018;21:81–89.

37. Chan AY, Urgun K, Tran DK, Kyong T, Hsu FPK, Vadera S. Cerebral Peduncle Volume and Motor Function Following Adult Hemispherectomy. World Neurosurg. 2019;126:156–159.

38. Küpper H, Kudernatsch M, Pieper T, et al. Predicting hand function after hemidisconnection. Brain. 2016;139:2456–2468.

39. Olulade OA, Seydell-Greenwald A, Chambers CE, et al. The neural basis of language development: Changes in lateralization over age. Proc Natl Acad Sci. 2020;117:23477–23483.

40. Ballantyne AO, Spilkin AM, Trauner DA. Language Outcome After Perinatal Stroke: Does Side Matter? Child Neuropsychol. 2007;13:494–509.

